# Biochemical characterization and mucosal immune function of IgT in grass carp (*Ctenopharyngodon idella*)

**DOI:** 10.1101/2025.05.04.652088

**Authors:** Bin Liu, Jing-Wei Zhao, Yong-An Zhang, Xu-Jie Zhang

**Author notes:** Correspondence: Yong-An Zhang,; Xu-Jie Zhang.

## Abstract

Immunoglobulin T (IgT) is a teleost-specific immunoglobulin discovered to be specialized in mucosal immunity. For fish that live in water throughout their lives, eliciting a local mucosal immune response through immersion immunization is an important and highly suitable way for preventing fish diseases. However, due to the lack of corresponding antibodies against IgT, studies on its immune responses to pathogens and vaccines in commercial fish have largely been limited to rainbow trout. Thus, a valuable tool for assessing mucosal immune response to immersion vaccination is still lacking in most commercial fish species. In this study, we expressed and purified recombinant protein of the heavy chain constant domains (CH) 1-2 of grass carp IgT and immunized mice to develop a mouse anti-grass carp IgT monoclonal antibody (mAb). Western blotting results showed that the mAb could specifically recognize grass carp IgT without cross-reacting with grass carp IgM, IgMT1 or IgMT2. We then measured IgM and IgT concentrations in the serum and gill mucus of grass carp and examined the responses of specific IgM and IgT to immersion immunization with grass carp reovirus genotype Ⅱ (GCRV-Ⅱ) and *Flavobacterium columnare*. We found that the concentration of IgT in gill mucus was higher than that in serum, and the IgT/IgM ratio in gill mucus was much higher than that in serum. Moreover, after immersion immunization, the specific IgT levels in gill mucus increased significantly, while no specific IgT was detected in serum. In contrast, the specific IgM levels in serum increased significantly, whereas there was little change in specific IgM levels in gill mucus. These results indicated that IgT is an important immunoglobulin in mucosal immune response, whereas IgM plays an important role in systemic immune response in grass carp. In summary, these findings not only further elucidate the biochemical and functional differences between IgM and IgT in grass carp, but also provide a valuable detection tool for accurately evaluating the immune efficacy of immersion vaccines in grass carp.

## 1. Introduction

The adaptive immune system is an important defense line that protects vertebrates from pathogenic invasion [1]. Immunoglobulins (Igs) are immune effector molecules secreted by B lymphocytes after activation and proliferation, and they play a crucial role in the recognition and binding of pathogens [2]. The basic structure of an immunoglobulin is a tetrapeptide composed of two heavy chains and two light chains linked by interchain disulfide bonds. Each heavy chain or light chain contains a constant region and a variable region [3]. Typically, the heavy chain constant (CH) region of an immunoglobulin is used to define its type of immunoglobulin. For example, in mammals, there are five immunoglobulin heavy chains: μ, δ, γ, α and ε, corresponding to the five immunoglobulin classes: IgM, IgD, IgG, IgA and IgE [4]. As primitive vertebrates, teleost fish possess an immune system similar to that of mammals. [5, 6]. To date, three immunoglobulin heavy chain types have been identified in teleost fish: μ, δ, and τ/ζ, which correspond to the immunoglobulin classes IgM, IgD [7], and IgT/IgZ [8, 9], respectively. Among them, IgM and IgD are conserved across vertebrates, while IgT is unique to teleost fish [6]. Since IgD mainly functions as a membrane receptor for B cells in vertebrates, and secreted IgD may play an evolutionary conserved role in mucosal homeostasis [10–12], the primary antibodies mediating humoral immune responses in teleost fish are IgM and IgT [6, 13].

In teleost fish, IgM is the most abundant immunoglobulin type, which is highly accumulated in both systemic and mucosal tissues and plays an important role in immune function [14, 15]. Before 2005, IgM was considered to be the only antibody type in teleost fish capable of responding to invading pathogens. However, this paradigm was challenged in 2005 with the discovery of a novel immunoglobulin gene, ighτ/ighζ, in zebrafish (*Danio rerio*) [8] and rainbow trout (*Oncorhynchus mykiss*) [9]. This gene encodes the heavy chain of the immunoglobulin IgT/IgZ (collectively referred to as IgT in this article). Since then, the antibody type of teleost fish has been redefined, and IgT is considered to be the last immunoglobulin type found in teleost fish [16]. To date, IgT has been identified in a variety of teleost fish, including rainbow trout [8, 17], zebrafish [8, 18], common carp (*Cyprinus carpio*) [19–21], grass carp (*Ctenopharyngodon idella*) [22], and others. Furthermore, several teleost species have been found to possess multiple IgT subclasses, such as two IgT subclasses in zebrafish [8, 18] and three IgT subclasses in rainbow trout [9, 17]. More interestingly, chimeric forms of IgT containing both Cμ and Cτ constant region domains have been identified in common carp [20], grass carp [22–24] and nile tilapia (*Oreochromis niloticus*) [25]. For example, grass carp possess two chimeric IgT: IgMT1 (with a CH region composed of “Cμ1-Cμ2-Cτ3-Cτ4”) and IgMT2 (with a CH region composed of “Cμ1-Cτ4”) [23, 24].

Functional studies of IgT have revealed it to be a specialized mucosal immunoglobulin that plays crucial roles in multiple mucosal surfaces, including the intestine [6], skin [26], gills [27], oropharyngeal cavity [28] and swim bladder [29, 30]. However, research on IgT-mediated immune responses to pathogens and vaccines in commercially important fish species has been largely restricted to rainbow trout, primarily due to the lack of species-specific anti-IgT antibodies. As a mucosa-specific immunoglobulin, the detection of IgT immune response is of great significance for the development of immersion vaccines.

Grass carp is one of the most important freshwater aquaculture species in China, accounting for a large proportion of total aquatic production [31]. In recent years, with the deterioration of aquaculture environment, grass carp diseases have occurred frequently. Vaccination represents the most effective and cost-saving ways to prevent fish diseases, and immersion vaccination is a necessary and suitable method for fish. Based on this, in the present study, we successfully developed a mouse anti-grass carp IgT mAb. Subsequently, we used the mouse anti-IgT mAb to identify the protein structures of the IgT in the serum, gill mucus and skin mucus of grass carp, and found that IgT existed as a monomer in serum, but as a polymer in gill mucus and skin mucus. And then, we determined the basal antibody concentrations of IgM and IgT in serum and gill mucus of grass carp, as well as the changes of specific IgM and IgT antibody levels in serum and gill mucus of grass carp after immersion immunization with *F. columnare* and GCRV-Ⅱ. The results showed that the IgT concentration in gill mucus was higher than that in serum. And the levels of specific IgT in gill mucus increased significantly after immersion immunization, while the serum specific IgT levels did not change. In contrast, serum specific IgM levels increased significantly, whereas specific IgM in gill mucus was almost unchanged. The results showed that IgT of grass carp, like IgT of rainbow trout, is an important mucosa-specific immunoglobulin, while IgM plays an important role in systemic immunity. These results not only enrich our understanding of functional differences between teleost fish IgM and IgT, indicating that teleost fish IgT is commonly specialized in mucosal immunity, but also provide a valuable tool for accurate evaluation of the immune effects of immersion vaccines in grass carp.

## 2. Materials and methods

### 2.1 Fish

Healthy grass carp (weighing 200 ± 20 g) were purchased from Huangpi Hatchery (Wuhan, Hubei, China) and maintained in laboratory aquarium tanks with water control system. Fish were fed with commercial pellets daily and acclimatized to the laboratory condition for at least 2 weeks before experiments.

### 2.2 Preparation and validation of mouse anti-grass carp IgT mAb

#### 2.2.1 Purification of antigen

The gene segment (GenBank accession NO. GQ201419.1) encoding the CH1-2 domains of grass carp IgT was amplified with primers IgT-CH1-2-F/R (shown in Table 1) and 2 × Taq Master Mix (Vazyme). The PCR product was digested with restriction enzymes *BamH* I and *Xhol* I (TAKARA), and ligated to the pET28a vector with an N-terminal SUMO tag, and then transformed into *E. coli* Rosetta (DE3). The recombinant protein SUMO-IgT-CH1-2 was expressed by induction with isopropyl *β*-D-1- Thiogalactopyranoside (IPTG, 1 mmol/L) at 18℃ for 12 h. The bacteria were collected by centrifugation (4℃, 5000 × *g*, 15 min), resuspended with PBS (pH = 7.4), and mechanically disrupted using a high-pressure homogenizer (ATS). The supernatant was collected by centrifugation (4℃, 17000 × *g*, 30 min) and filtered through a 0.22 μm filter, and then passed through a Ni column. The column was next washed sequentially with PBS containing 20 mM, 60 mM, 200 mM, and 500 mM imidazole. The target protein was identified by sodium dodecyl sulfate polyacrylamide gel electrophoresis (SDS-PAGE) and the imidazole was removed by dialysis. Finally, the protein concentration was detected by measuring the absorption at 562 nm by BCA protein assay kit (Biosharp) with BSA as a standard protein.

**Table 1.**
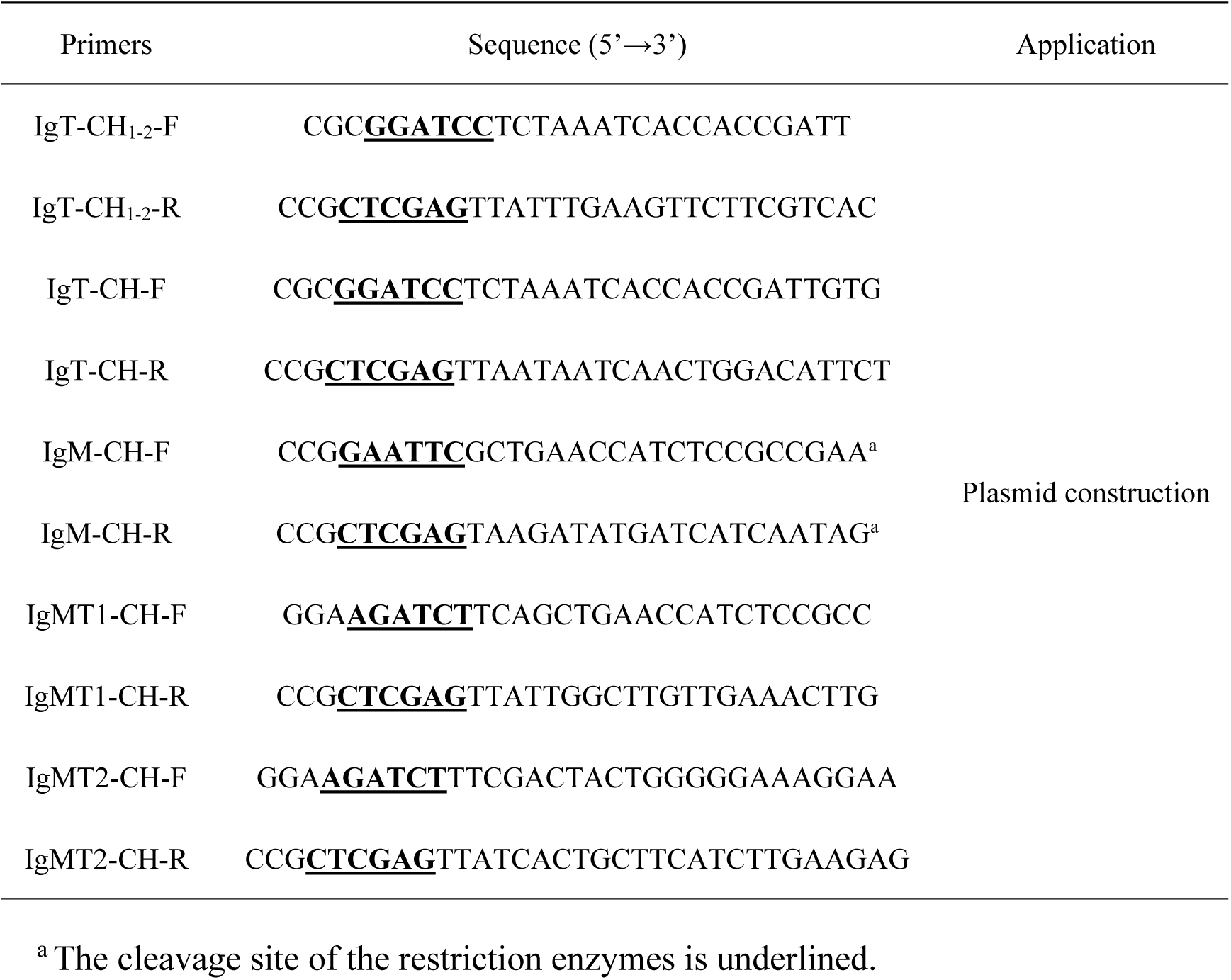
Primers used in this study.

#### 2.2.2 Mice immunization and hybridoma cell preparation

The recombinant protein SUMO-IgT-CH1-2 was applied to immunize mice to raise the mAb (CUSABIO Laboratories). Briefly, six BALB/c female mice (18–20 g) were immunized three times with 50 μg per dose of the immunogen at 14-day intervals. In the first immunization Freund’s complete adjuvant (FCA) (Sigma-Aldrich) was used, while Freund’s incomplete adjuvant (FIA) (Sigma-Aldrich) was used in the second and third immunization. Spleen cells from the inoculated mice were fused with myeloma cells to prepare hybridoma cells. Briefly, spleen cells from immunized mice were mixed with myeloma cells (SP2/0-Ag 14, ATCC) at a ratio of 5:1, then centrifuged at 1000 rpm for 10 min and the supernatant was discarded. Then, 1 mL of PEG-1450 was slowly added to the tube with gentle shaking. Subsequently, 40 mL of Dulbecco’s modified eagle’s medium (DMEM) was gradually added in multiple steps. The entire procedure was performed in a 37℃ water bath. After fusion, the mixture was centrifuged at 1,000 rpm for 10 min, the supernatant was carefully removed, and the pellet was resuspended in hypoxanthine-aminopterin-thymidine (HAT) medium supplemented with 20% fetal bovine serum (FBS) to obtain the fused cells.

#### 2.2.3 Screening of hybridoma cells and preparation of antibodies

The supernatant of hybridoma cells exhibiting specific reactivity with IgT-CH1-2 was screened by two independent enzyme-linked immunosorbent assay (ELISA). Briefly, the recombinant protein SUMO-IgT-CH1-2 or SUMO (2 μg/mL diluted in 0.05 M carbonate buffer; 100 μL/well) were added to 96-well plate (BioFil). After coating overnight, the plate was washed 5 times with PBST (PBS containing 0.05% Tween-20). Next, skim milk (5% in PBST, 250 μL) was added to each well, and the plate was incubated at 37℃ for 2 h and washed. Then, the supernatant of hybridoma cells (100 μL) was added to each well, followed by incubation at 37℃ for 2 h. Then HRP-conjugated goat anti-mouse IgG Abs (Proteintech) (1:5000 diluted in PBST) were added (100 μL/well) and incubated at 37℃ for 1 h. Finally, TMB chromogen solution (100 μL, Biosharp) was added to each well and incubated for 30 min, and then 2 M H2SO4 (50 μL/well) was added for 10 min to stop the reaction. The absorbance was measured with a microplate reader (Bio-Tek) at 450 nm. The cells showing strong reactivity with SUMO-IgT-CH1-2 but not SUMO were selected as the final hybridoma cells. Finally, one positive clone (IgT-6H6G11) from the two candidates (IgT-6H6G11 and IgT-2H2H2) was injected intraperitoneally into mice, and ascites was harvested after 14 days. The IgG fraction from the ascites was purified using a HiTrap Protein A+G agarose column (GE Healthcare) according to the instructions of the manufacturer.

#### 2.2.4 Determination of anti-IgT mAb subclass

Determination of mAb subclass of selected hybridoma cell was performed by an ELISA assay under the same conditions as described above, except for the secondary antibody used. In this case, 100 μL of the recombinant protein SUMO-IgT-CH1-2 was coated to 96-well plate and incubated at 4℃ overnight. After blocking by 5% skim milk at 37℃ for 2 h, the selected hybridoma supernatant was added to each well and incubated at 37℃ for 2 h. And then HRP conjugated pAbs against mouse IgG1, IgG2a, IgG2b, IgG2c, IgG3, IgA or IgM were added to the corresponding wells and incubated at 37℃ for 1 h. Then, TMB chromogen solution was added to each well and incubated for 30 min, followed by the addition of 2 M H2SO4 to stop the reaction after 10 min. Absorbance was measured with a microplate reader at 450 nm and the subclass of antibody was determined according to the positive reactivity observed.

#### 2.2.5 Cloning and expression of grass carp IgT, IgM, IgMT1, and IgMT2 heavy chain constant region

The gene segments encoding the constant regions of grass carp IgT, IgM, IgMT1, and IgMT2 (GenBank accession nos. GQ201419.1, DQ417927.1, GQ201421.1 and GQ201420.1, respectively) were amplified using the corresponding primers (shown in Table 1) and the PCR products were digested with appropriate restriction enzymes and ligated to the pGEX-6p-1 vector with an N-terminal GST tag, and then transformed into *E. coli* BL21 (DE3). Recombinant protein expression was induced with IPTG at 1 mmol/L.

#### 2.2.6 Western blotting

The specificity of the mouse mAb was evaluated by Western blotting. The recombinant proteins of grass carp IgT, IgM, IgMT1, and IgMT2 heavy chain constant regions were resolved on 8-12% gradient polyacrylamide gels and transferred to PVDF membrane. The membrane was blocked for 2 h at room temperature in TBST buffer (25 mM TrisHCl, 150 mM NaCl, 0.1% Tween 20, pH 7.5) containing 5% skim milk powder, followed by incubation with mouse anti-grass carp IgT mAb or rabbit anti-GST pAb (2 μg/mL diluted in TBST) at 4℃ overnight. After washing three times with TBST, the membrane was incubated with HRP-conjugated anti-mouse IgG or HRP-conjugated anti-rabbit IgG (Proteintech; 1:5000 dilution in TBST) for 1 h at 37℃. After additional three times washing with TBST, the membranes were incubated with Clarity Western ECL Substrate (Abbinke) and imaged using the Amersham Imager 600 (GE Healthcare).

### 2.3 Collection of serum, gill mucus and skin mucus

For sampling, grass carp were anesthetized with MS-222 (Sigma). Peripheral blood was collected from the tail vein and allowed to clot at 4℃ for 4 h, and then centrifuged to obtain serum. Gill mucus collection was performed as previously described [27]. Briefly, blood in the gills was first removed by cardiac infusion of PBS-heparin until the gills were completely bleached. The branchial arches were removed and the gill lamellae were washed three times with PBS to eliminate residual blood. After that, gills were incubated with protease inhibitor buffer (1 × PBS containing 1 × protease inhibitor cocktail (Roche), 1mM phenylmethylsulfonyl fluoride (Sigma); Ph 7.2) (1 mL buffer per 1 g) for 12 h. And the fluid was then collected and centrifuged at 400 × *g* for 10 min at 4℃ to obtain the supernatant. Grass carp skin mucus was collected as described [26]. Briefly, the mucus was gently scraped from the skin surface, transferred into an Eppendorf tube, vigorously vortexed, and centrifuged at 400 × *g* for 10 min at 4℃ to obtain the supernatant.

### 2.4 Gel filtration

To analyze the monomeric or polymeric state of IgT and IgM in grass carp serum, gill mucus and skin mucus, gel filtration analyses were performed using a Superdex^TM^ 200 Increase 10/300 GL (GE Healthcare) as previously described [6]. IgM and IgT in the eluted fractions were identified by Western blot analysis and ELISA using IgM- and IgT-specific mAbs, respectively.

### 2.5 Detection of IgM and IgT concentrations in serum and gill mucus of grass carp

The total IgM and IgT concentration in grass carp serum and gill mucus were determined by ELISA as described by Velázquez et al. [32]. Briefly, the serum (diluted 1:100) and gill mucus (diluted 1:5) in 0.05 M carbonate buffer was added to 96-well plate (100 μL/well; BioFil). After coating overnight at 4℃, the plate was washed 5 times with PBST. The wells were then blocked with 5% skim milk (250 μL/well) was at 37℃ for 2 h, followed by washing. Then, mouse anti-grass carp IgM or IgT mAbs (2 μg/mL) were added to respective well and incubated at 37℃ for 1.5 h. After that, HRP-conjugated goat anti-mouse IgG Abs (Proteintech) were subsequently added and incubated at 37℃ for 1 h. Finally, TMB chromogen solution (100 μL, Biosharp) was added to each well and incubated for 30 min, and then 2 M H2SO4 (50 μL) was added for 10 min to stop the reaction. Absorbance was measured at 450 nm using a microplate reader (Bio-Tek). For quantitative analysis of serum and gill mucus total IgM and IgT concentration, standard curves were generated using serial dilutions (from 0.015625 to 0.25 μmol/L) of recombinant IgM or IgT heavy chain constant regions.

### 2.6 Immunization and sampling of grass carp

To explore whether anti-grass carp IgT mAb could be used to detect specific IgT and elucidate the function of IgT, grass carp were immunized with bacterial and viral pathogens, followed by monitoring of IgM and IgT responses in serum and gill mucus. The bacteria used in this study was *F. columnare* G4 strain stored in our laboratory. The strain was streaked from −80°C freezer and inoculated into Shieh solid medium and cultured at 28℃ for 16 h. Then the monoclonal strain was selected and transferred to Shieh liquid medium at 28℃ for further culture, and the virus used for infection in this study was GCRV-Ⅱ (GCRV106). For immunization, 60 fish (50 ± 5 g) were bath-exposed with *F. columnare* (5×10^8^ CFU) or GCRV-Ⅱ (1 × 10^8^ copies) in 10 L water for 4 h at 28℃. For GCRV-Ⅱ immunization, a second immersion vaccination was performed at 14 days after the primary vaccination. Samples including serum and gill mucus were collected from 4 individuals at days 0, 3, 7, 10, 14 and 21 after vaccination. Throughout this time, the fish were maintained in a flow through aquaria at 28℃, and fed daily with dry pellets at 0.5-1% biomass.

### 2.7 Analysis of IgM and IgT response levels

The specific IgM and IgT in serum and gill mucus of grass carp immunized with *F. columnare* and GCRV-Ⅱ were detected by ELISA. Briefly, inactivated *F. columnare* (1 × 10^9^ CFU/mL; 100 μL/well) or GCRV-Ⅱ (5 × 10^3^ copies/μL; 100 μL/well) were coated on 96-well plates overnight at 4℃. Next, skim milk (5%, 250 μL) was added to each well, and the plate was incubated at 37℃ for 2 h and washed. Then, the serum (1:100 dilution) and gill mucus (1:5 dilution) in PBS was added and incubated at 37℃ for 3 h. Subsequently, mouse anti-grass carp IgM or IgT mAb (1 μg/mL) was added to each well and incubated at 37℃ for 1 h. After that, HRP-conjugated goat anti-mouse IgG Abs (Proteintech) were added and incubated at 37℃ for 1 h. Finally, TMB chromogen solution (100 μL, Biosharp) was added to each well and incubated for 30 min, and then 2 M H2SO4 (50 μL) was added for 10 min to stop the reaction. The absorbance was measured with a microplate reader (Bio-Tek) at 450 nm.

### 2.8 Statistical analysis

The statistic *p*-value was calculated using one-way ANOVA with a Dunnett’s *post hoc* test (SPSS Statistics, Version 19, IBM). The *p*-values lower than 0.05 were considered statistically significant.

## 3. Results

### 3.1 Production and characterization of mAb against grass carp IgT

There are three subclasses of grass carp IgT, including IgT (the heavy chain constant region is composed of “Cτ1-Cτ2-Cτ3-Cτ4”), IgMT1 (the heavy chain constant region is composed of “Cμ1-Cμ2-Cτ3-Cτ4”), and IgMT2 (the heavy chain constant region is composed of “Cμ1-Cτ4”). To prepare mAb that specifically recognize IgT but not the other two subclasses, the CH1-2 domains of IgT (Cτ1-Cτ2) with low similarity to the constant domains of other immunoglobulins were selected for fusion expression with an N-terminal SUMO tag (SUMO-IgT-CH1-2, Fig 1A). After a series of purification procedures, a highly purified recombinant protein was obtained (Figure 1B, purity >90%) and used to immunize mice to generate hybridoma cells. To screen hybridoma cell lines producing antibodies specific to grass carp IgT, positive clones were first screened by ELISA using the recombinant protein SUMO-IgT CH1-2 as the coating antigen, followed by reverse screening with purified recombinant protein SUMO (FIG. 1C). Finally, a clone (6H6G11, IgG2a isotype) that positively recognized SUMO-IgT CH1-2 but negatively recognized SUMO was selected as the final cell line for preservation and antibody preparation (Fig 1D).

**FIGURE 1.**
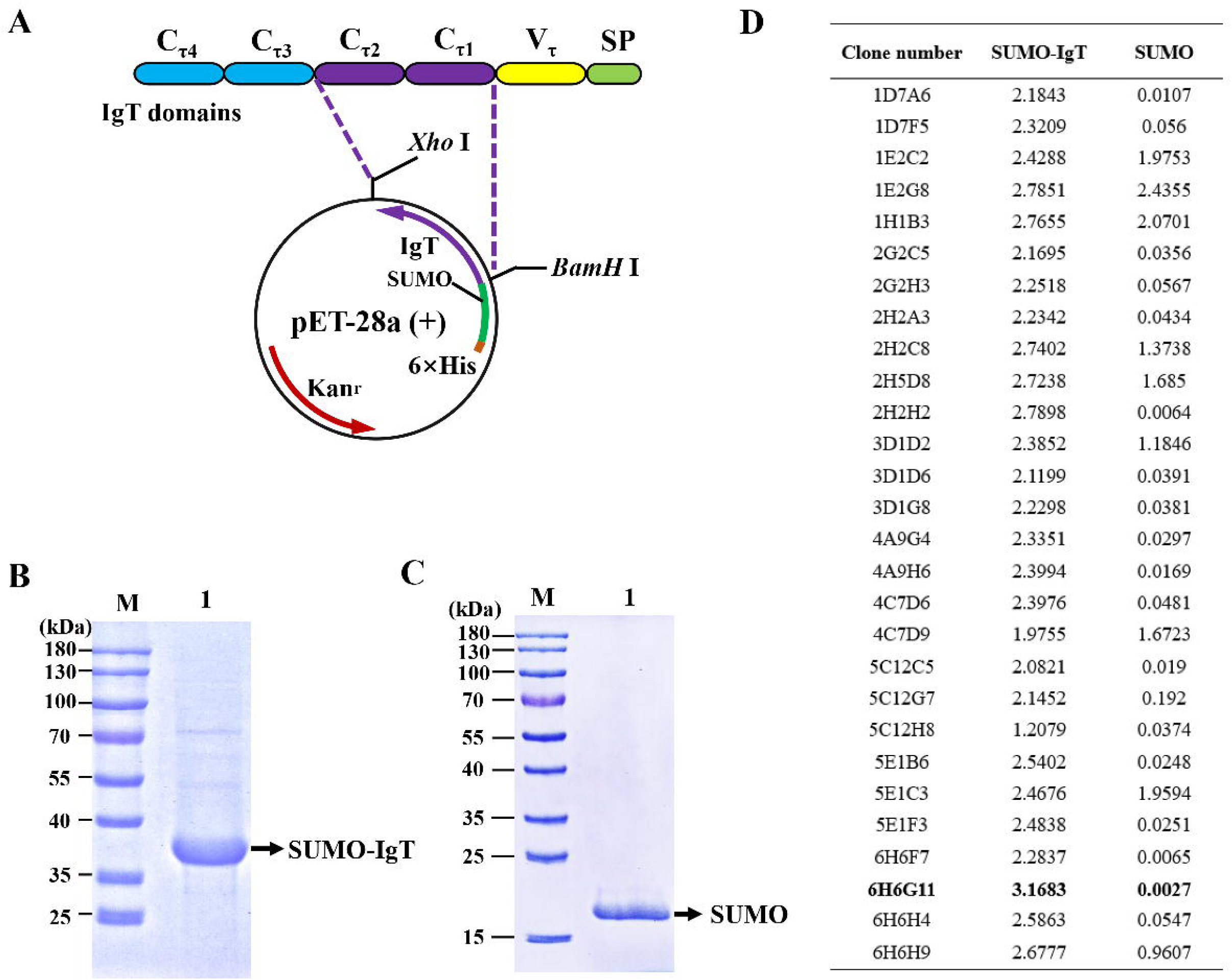
Development of monoclonal antibodies against grass carp IgT. (A) The expression plasmid to produce a recombinant fragment of grass carp IgT was constructed by inserting the cDNA encoding for the CH1-2 domains of grass carp IgT heavy chain into a modified pET28a (+) vector with an N-terminal SUMO-6×His-Tag. SP, signal peptide; Vτ, variable domain of grass carp IgT heavy chain; Cτ1-4, constant domains 1-4 of grass carp IgT heavy chain. (B) The purified recombinant protein SUMO-IgT CH1-2 fragment (∼2 μg) was resolved on a 12% SDS-PAGE and stained with Coomassie blue. Lane M, protein marker; Lane 1: recombinant SUMO-IgT CH1-2 fragment. (C) The purified recombinant protein SUMO (∼2 μg) was resolved on a 15% SDS-PAGE and stained with Coomassie blue. Lane M, protein marker; Lane 1: SUMO. (D) The selected hybridoma cell line clone 6H6G11 recognized IgT but not SUMO.

### 3.2 Specificity analysis of IgT mAb

To verify the specificity of the mouse anti-grass carp IgT mAb, we tested it at recombinant protein level. First, we constructed recombinant plasmids containing the heavy chain constant regions of grass carp IgM, IgT, IgMT1 and IgMT2. The recombinant proteins including GST-IgM CH (Fig 2A), GST-IgT CH (Fig 2B), GST- IgMT1 CH (Fig 2C) and GST-IgMT2 CH (Fig 2D) were expressed by *E. coli* and subjected to Western blotting analysis. The results demonstrated that the mouse anti-grass carp IgT mAb specifically recognize grass carp IgT but showed no cross-reactivity with grass carp IgM (Fig 2E) or with the other IgT subclasses, IgMT1 and IgMT2 (Fig 2F). These results showed that the antibody was specific for grass carp IgT.

**FIGURE 2.**
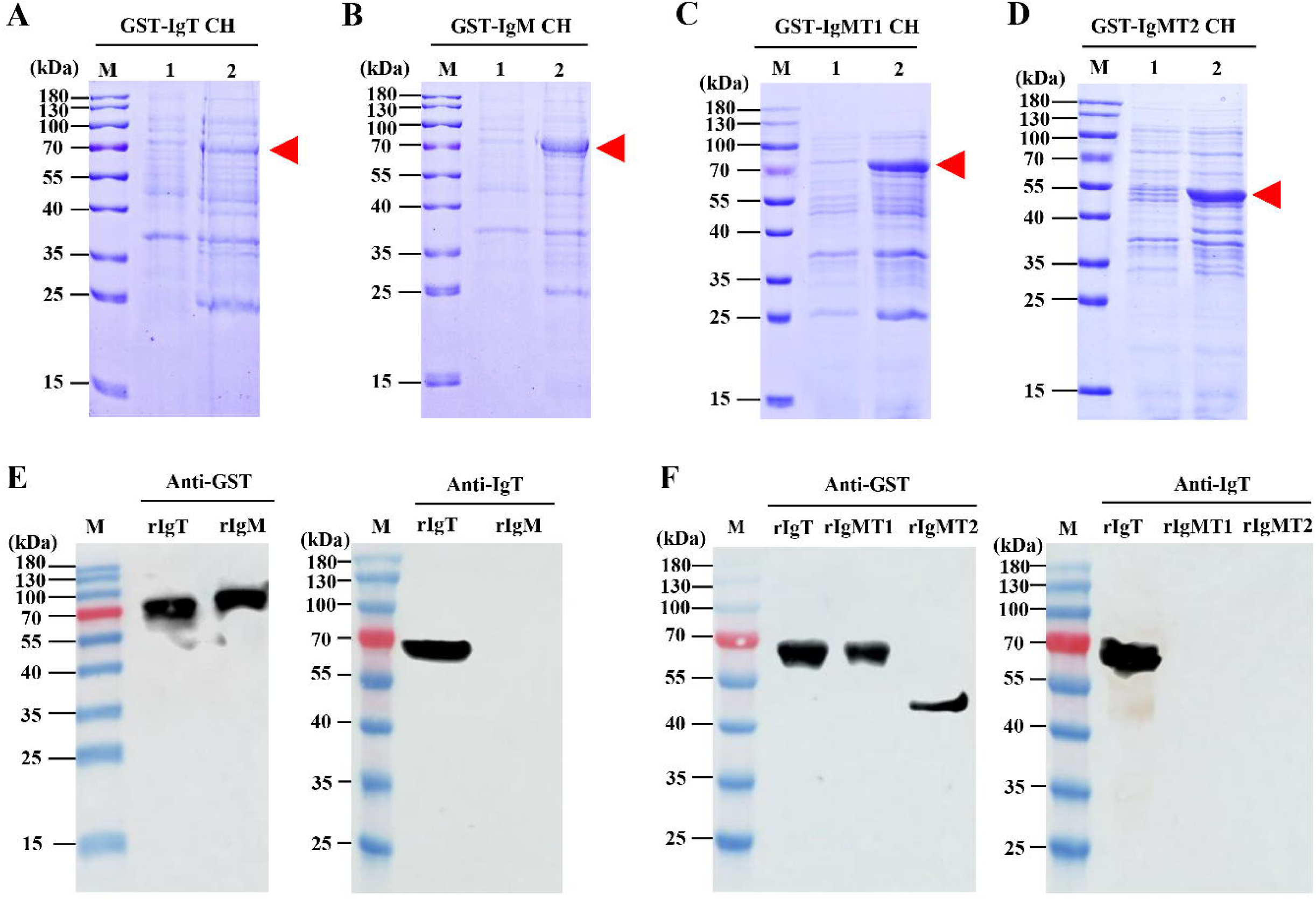
Specificity analysis of mouse anti-grass carp IgT mAb with SDS-PAGE and Western blotting assessment. (A-D) Prokaryotically expressed GST-IgT CH (A), GST-IgM CH (B), GST-IgMT1 CH (C) and GST-IgMT2 CH (D) were resolved on a 12% SDS-PAGE and stained with Coomassie blue. Red arrowheads indicate the corresponding target band. M, protein marker; Lane 1, total cellular extracts from *E. coli* BL21 (DE3) containing expression vector without IPTG induction; Lane 2, total cellular extracts from *E. coli* BL21 (DE3) containing expression vector with IPTG induction. (E) Cellular extracts containing IgT CH and IgM CH were immuno-stained with anti-GST or anti-IgT mAb. M, Protein marker; rIgT and rIgM: total cellular extracts from *E. coli* BL21 (DE3) containing expressed GST-IgT CH and GST-IgM CH with IPTG induction, respectively. (F) Cellular extracts containing IgT CH, IgMT1 CH and IgMT2 CH were immuno-stained with anti-GST or anti-IgT mAb. M, protein marker; rIgT, rIgMT1 and IgMT2: total cellular extracts from *E. coli* BL21 (DE3) containing expressed of GST-IgT CH, GST-IgMT1 CH and GST-IgMT2 CH with IPTG induction, respectively.

### 3.3 Protein characterization of serum, gill mucus and skin mucus IgT

So far, studies on IgT structure have been limited to rainbow trout and no related studies in other fish species. To characterize the structure of IgT in grass carp, we collected serum, gill mucus and skin mucus and preliminarily purified by gel filtration, respectively. Eluted fractions were detected by Western blotting and ELISA. The results showed that serum IgT (Fig 3A) eluted exclusively at the position expected for a monomer (∼180 kDa) based on the standard curve generated by gel filtration molecular weight standards. In contrast, most of the IgT in gill mucus (Fig 3B) and skin mucus (Fig 3C) was polymeric, as it was eluted at a position similar to grass carp IgM, a tetrameric immunoglobulin. Subsequently, the samples with elution volumes of 9.5 mL and 13.5 mL were subjected to Western blotting under non-reducing conditions, and the results further confirmed that serum IgT was approximately 180 kDa, while serum IgM was over 300 kDa (Fig 3D). Conversely, IgT in gill mucus (Fig 3E) and skin mucus (Fig 3F) exhibited a molecular weight similar to that of IgM, indicating a polymeric form. ELISA results were consistent with the Western blotting data and further demonstrated that serum IgT exists a monomer (Fig 3G), whereas IgT in gill mucus (Fig 3H) and skin mucus (Fig 3I) forms polymers similar to IgM.

**FIGURE 3.**
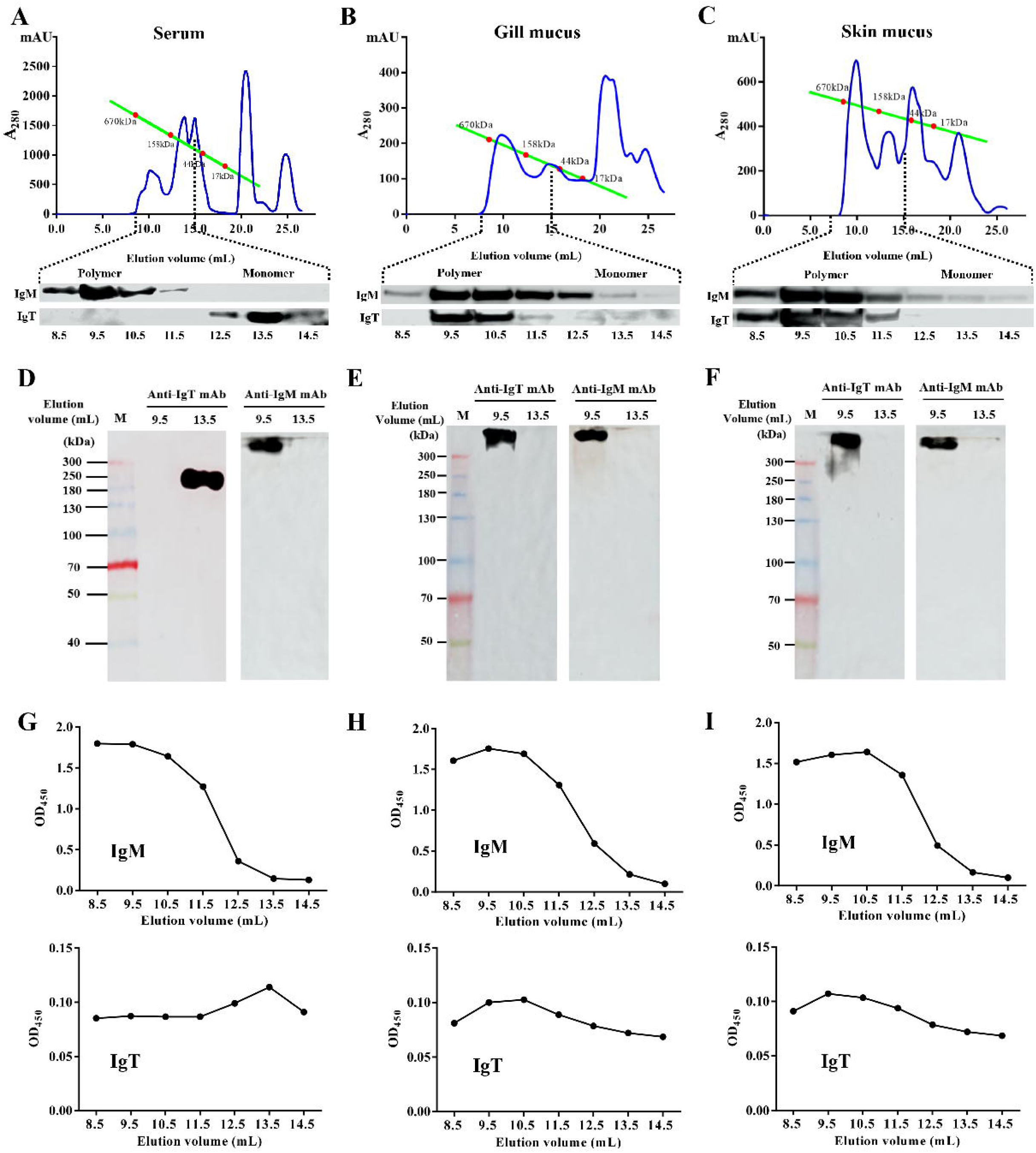
Structural characterization of IgT. (A-C) Fractionation of serum (A), gill mucus (B) or skin mucus (C) (top), followed by immunoblot analysis of the fractions with IgM and IgT mAb (below). *A280*, absorbance at 280 nm. (D-F) 10% SDS-PAGE of gel-filtration fractions of serum (D), gill mucus (E) or skin mucus (F) corresponding to elution volumes of 9.5 mL and 13.5 mL under nonreducing conditions, followed by immunoblot analysis with mAb to grass carp IgM or IgT. (G-I) ELISA detection of gel-filtration fractions of serum (G), gill mucus (H) or skin mucus (I) corresponding to elution volumes from 8-15 mL with mAb to grass carp IgM (top) or IgT (below).

### 3.4 Analysis of IgM and IgT concentrations in serum and gill mucus of grass carp

Previous studies have shown that specialization of immunoglobulin isotypes into mucosal and systemic responses occurred during tetrapod evolution [27]. Based on this, to explore the immune functions of grass carp IgM and IgT in the systemic or mucosal regions, we quantified the concentrations of the IgT and IgM in serum and gill mucus of grass carp. First, we purified recombinant proteins of the IgT and IgM constant regions (Fig 4A) and used them to generate standard curves for IgT (Fig 4B) and IgM (Fig 4C). And then we analyzed the protein concentrations of IgT and IgM in serum and gill mucus and found that the IgM concentration in serum (about 2788 μg/mL) was much higher than in gill mucus (about 46.7 μg/mL). In contrast, the IgT concentration in serum (about 7.7 μg/mL) was slightly lower than in gill mucus (about 7.9 μg/mL) (Fig. 4 D and E). Additionally, the ratio of IgT to IgM was 62-fold higher in the gill mucus than in serum (Fig. 4F), suggesting that IgT might play a critical role in gill mucosal immunity.

**FIGURE 4.**
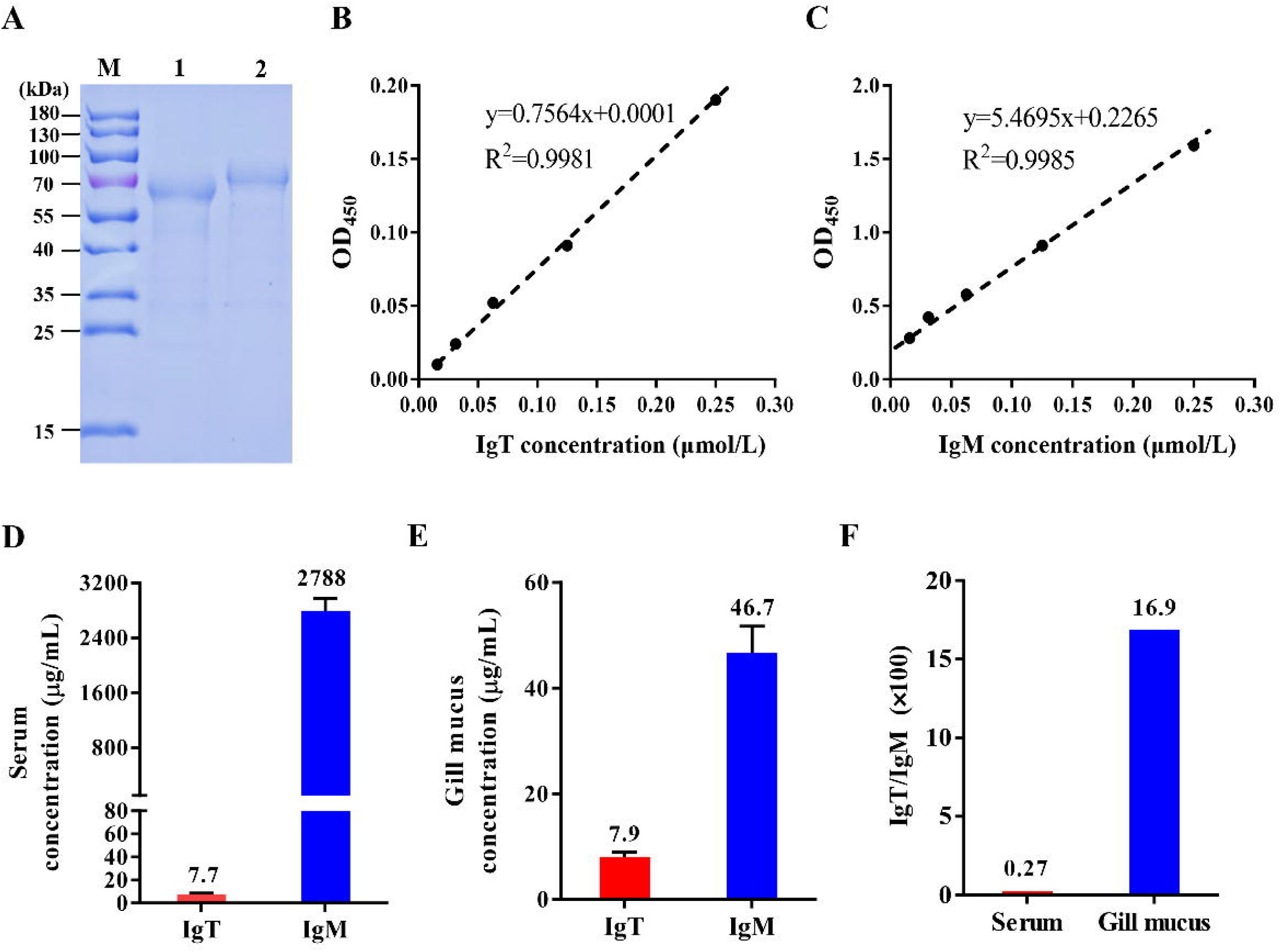
Detection of IgM and IgT basic antibody concentrations in serum and gill mucus. (A) Purified grass carp IgT CH and IgM CH recombinant proteins were resolved on a 12% SDS-PAGE and stained with Coomassie blue. Lane M, protein marker; Lane 1: recombinant GST-IgT-CH; Lane 2: recombinant GST-IgM-CH. (B-C) The standard curve constructed using the recombinant protein of grass carp IgT (B) and IgM (C) heavy chain constant region. (D-E) ELISA analysis of the concentration of IgM and IgT in serum (D) and gill mucus (E); n = 5 fish. (F) Ratio of IgT to IgM in serum and gill mucus, calculated from the values in D and E. Data are representative of at least three independent experiments.

### 3.5 Analysis of specific IgM and IgT levels in serum and gill mucus after GCRV-Ⅱ immunization

GCRV-Ⅱ is the pathogen of grass carp hemorrhagic disease, causes significant economic losses in grass carp cultivation annually. It is of great significance to develop mAbs to accurately evaluate the immune effects of vaccines for promoting the development of GCRV-Ⅱ vaccines. To investigate whether anti-IgT mAb can be used to detect specific IgT levels in grass carp, here, GCRV-Ⅱ was used to immunize grass carp by immersion, and the immune response levels of IgM and IgT in serum and gill mucus after immunization were analyzed. The results demonstrated that GCRV-Ⅱ immunization induced strong immune response in grass carp. The serum specific IgM levels continued to increase from day 10 after immunization (Fig. 5A), whereas the levels of specific IgM in gill mucus increased only on day 14 and then decreased immediately (Fig. 5B). In contrast, there was little change in specific IgT levels in serum after immunization (Fig. 5C), whereas specific IgT levels were persistently elevated in gill mucus (Fig. 5D). These results suggested that IgT, as a mucosa-specific immunoglobulin, plays an important role in mucosal immunity of grass carp after virus immunization, while IgM mainly plays a role in systemic immunity. And the results confirmed that anti-IgT mAb is an important and effective tool for the detection of mucosal immune response after virus immunization.

**FIGURE 5.**
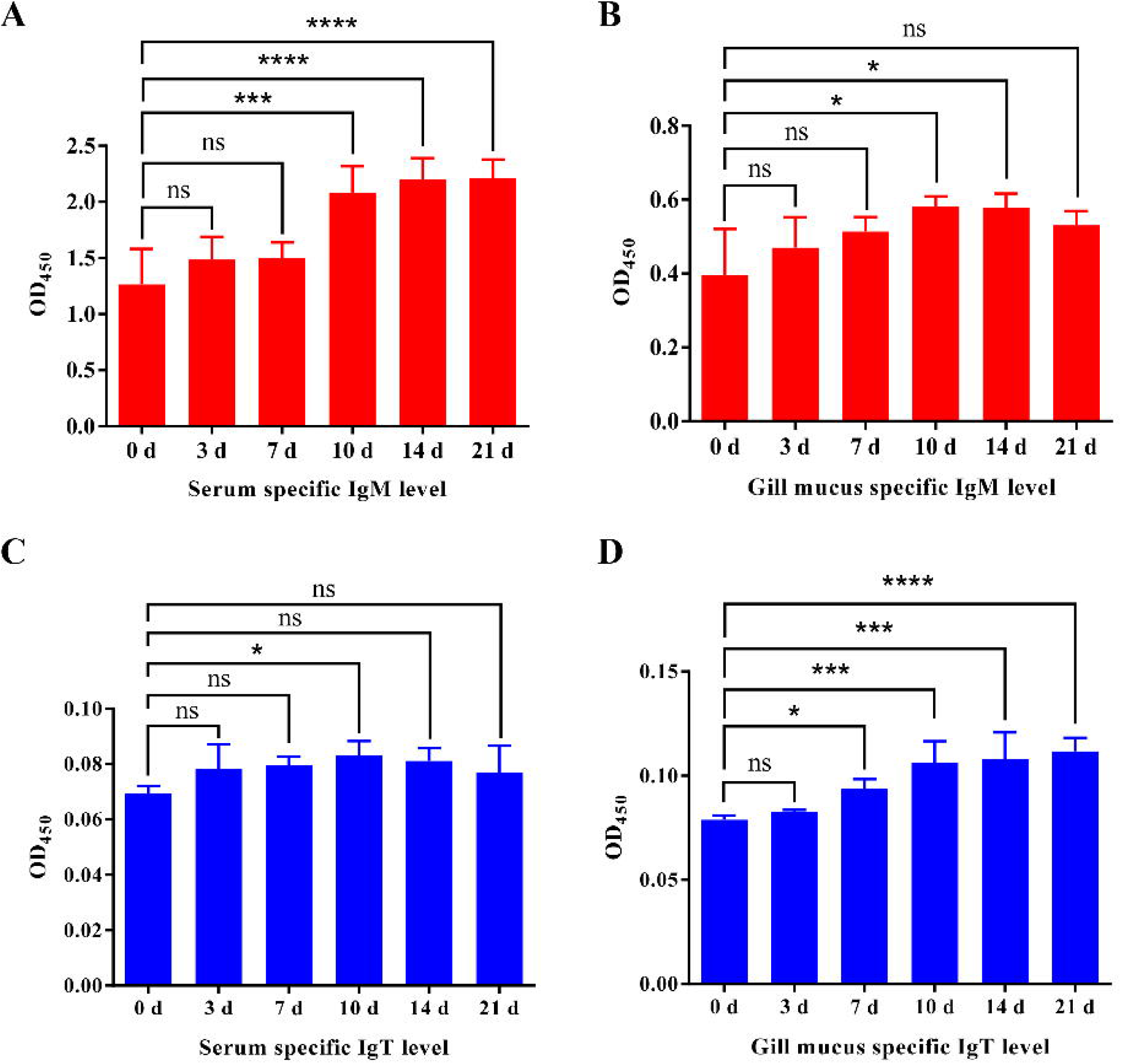
Analysis of IgM and IgT responses in serum and gill mucus after GCRV- Ⅱ immunization. (A) The changes of serum specific IgM levels after immunization. (B) The changes of gill mucus specific IgM levels after immunization. (C) The changes of serum specific IgT levels after immunization. (D) The changes of gill mucus specific IgT levels after immunization. Data shown are the mean ± SEM (n = 4 fish). The statistic *p* value was calculated by one-way ANOVA with a Dunnett *post hoc* test (**\****p* < 0.05, **\*\*\****p* < 0.001, **\*\*\*\****p* < 0.0001)

### 3.6 Analysis of specific IgM and IgT levels in serum and gill mucus after *F. columnare* immunization

*F. columnare* is a bacterial pathogen that seriously harms freshwater fish. It mainly infects the gills, which are important mucosal immune and respiratory organs in fish. To evaluate the immune response of grass carp gill mucosa after immunization with *F. columnare*, we immunized grass carp by immersion with *F. columnare* and detected IgM and IgT responses in the serum and gill mucus after immunization. The results showed that in the serum samples, specific IgM levels increased significantly on days 7, 10, 14, and 21 post-immunization (Fig 6A), whereas specific IgT levels only showed a slight increased on day 10 and subsequently became nearly undetectable (Fig 6C). On the contrary, in the gill mucus samples, specific IgT levels began to rise significantly on day 7 post-immunization, with a more pronounced increase from day 14 onward. Specific IgM levels in gill mucus also increased, but the response was less prominent compared to IgT (Fig 6 B and D). These results indicated that *F. columnare* immunization induced a strong IgT response in the gill mucosa, indicating that the anti-IgT mAb is a valuable tool for detecting gill mucosal immune responses after bacterial immunization.

**FIGURE 6.**
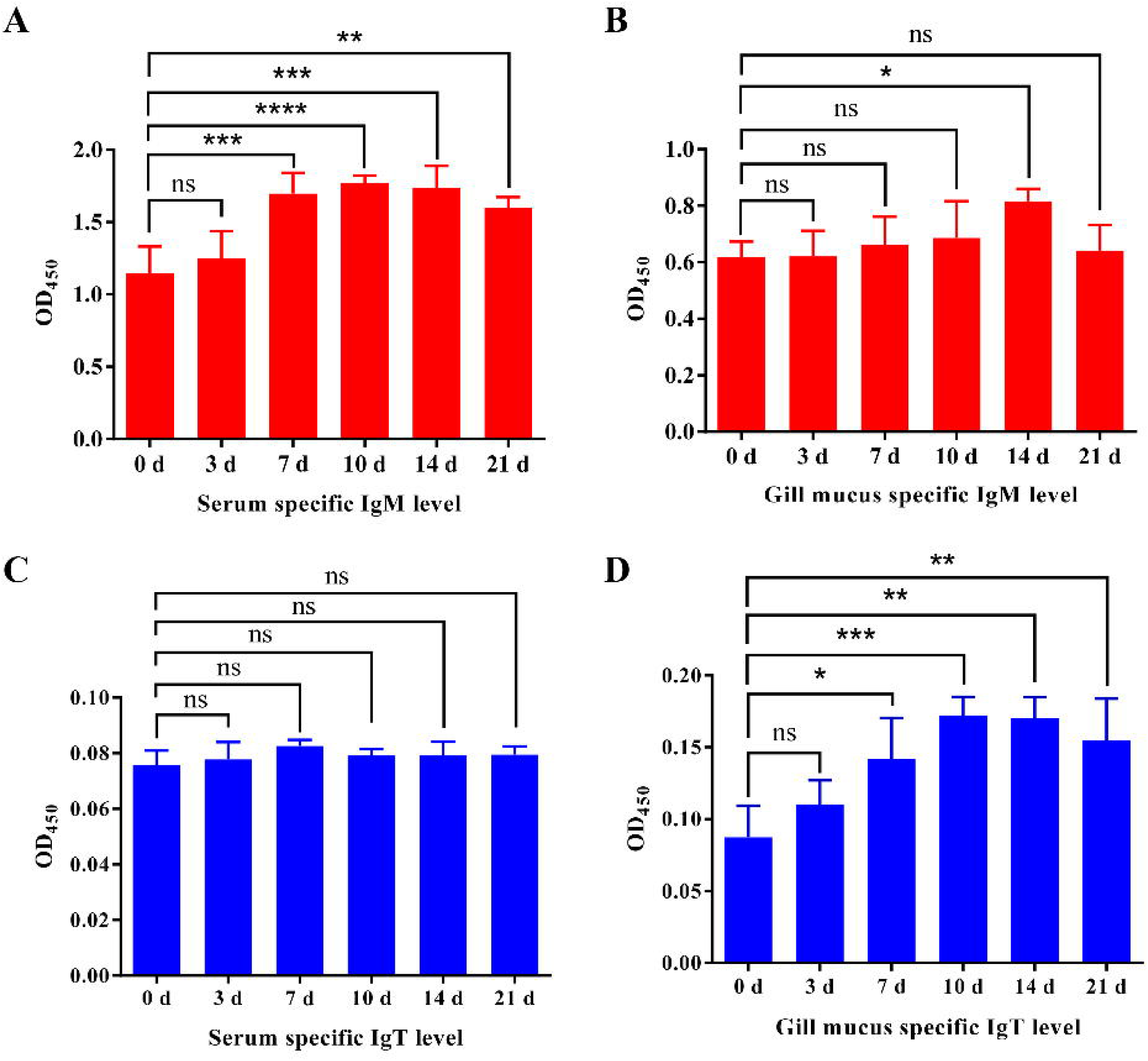
Analysis of IgM and IgT responses in serum and gill mucus after *F. columnare* immunization. (A) The changes of serum specific IgM levels after immunization. (B) The changes of gill mucus specific IgM levels after immunization. (C) The changes of serum specific IgT levels after immunization. (D) The changes of gill mucus specific IgT levels after immunization. Data shown are the mean ± SEM (n = 4 fish). The statistic *p* value was calculated by one-way ANOVA with a Dunnett *post hoc* test (**\****p* < 0.05, **\*\****p* < 0.01, **\*\*\****p* < 0.001, **\*\*\*\****p* < 0.0001)

## 3. Discussion

The discovery of IgT in teleost fish in 2005 attracted significant attention, as it challenged the existing understanding of the immune system of teleost fish. As the most recently identified and the last class of immunoglobulins found in teleost fish, the function and biochemical characteristics of IgT remained unknown until 2010. Following the successful preparation of mouse anti-rainbow trout IgT mAb, the function of rainbow trout IgT was revealed in 2010. These studies demonstrated that IgT functions similarly to IgA in mammals and birds [33], serving as an immunoglobulin specialized in mucosal immunity [6]. However, due to the lack of specific antibodies against IgT, studies on its immune responses to pathogens and vaccines in commercial fish have largely been limited to rainbow trout.

In this study, the CH1-2 domains of grass carp IgT were prokaryotically expressed and purified, and mouse anti-grass carp IgT mAb was successfully developed. It has been proved that the mAb could specifically recognize the heavy chain of grass carp IgT without cross-reacting with other grass carp immunoglobulins, including the other two subclasses of grass carp IgT. Notably, grass carp possess three IgT subclasses (IgT, IgMT1 and IgMT2), and it is very difficult to prepare monoclonal antibodies that recognize only one IgT subclass. So far, antibodies specifically recognizing a certain subclass of IgT have been reported in only a few teleost species [34]. The successful preparation of mAb targeting IgT subclasses in this study is of great significance for investigating the function of IgT subclasses and evaluating the immune efficacy of fish vaccines. However, after extensive validation, the anti-grass carp IgT mAb developed in this study was found to be suitable for Western blot and ELISA detection of IgT, but not for flow cytometric identification of IgT^+^ B cells.

Previous studies have shown that the polymerization state of IgT in rainbow trout serum is quite different from that in mucus, where IgT exists as a monomer in serum but as a polymer in mucus [6]. To explore the polymerization state of IgT in grass carp, we used mouse anti-IgT mAb for detection and found that, consistent with observations in rainbow trout, grass carp IgT exists as a monomer in serum but as a polymer in gill mucus and skin mucus. However, unlike rainbow trout, where polymeric IgT dissociates into monomers under non-reducing conditions in SDS-PAGE, the mucosal polymeric IgT in grass carp does not dissociate under the same conditions. These results suggested that the structure of polymeric IgT in grass carp is more stable than that in rainbow trout or that the monomer subunits of polymerized IgT in grass carp are not associated through non-covalent interactions like those in rainbow trout. These findings highlight potential interspecies differences in the structural stability or polymerization mechanisms of teleost polymeric IgT. Subsequently, we measured the concentrations of IgM and IgT in serum and gill mucus of grass carp. Similar to rainbow trout, the IgT/IgM in gill mucus was significantly higher than that in serum, indicating that IgT may play an important role in gill mucus.

As aquatic organisms, fish are constantly exposed to a pathogen-rich environment, making the mucosal immune tissues their first line of defense against infection. Vaccination can activate the immune system of fish and enhance resistance to specific pathogens, representing a critical strategy for disease prevention in commercial fish [35]. At the same time, accurate evaluation of vaccine immune effect is a very important step in vaccine development. To determine whether the anti-grass carp IgT mAb prepared in this study can be used to assess the immune efficacy of immersion vaccine, the most suitable for fish vaccination, grass carp were immunized via immersion with GCRV-Ⅱ and *F. columnare*, the causative agents of two major grass carp diseases. The levels of antigen-specific IgM and IgT in serum and gill mucus were then measured. The results showed that IgT exhibited a strong mucosal immune response in gill mucosa after immunization with either viral or bacterial antigens, while specific IgM levels increased significantly in serum. These findings align with observations in rainbow trout and not only confirm the importance of IgT in gill mucosal immunity but also provide a valuable tool for grass carp vaccine development and immune efficacy evaluation.

In conclusion, this study elucidated the biochemical characteristics of IgT in grass carp and its key role in the mucosal immunity, providing more further evidence that teleost fish IgT is commonly specialized in mucosal immunity. Additionally, it offers an important tool for advancing research on grass carp immune system and for disease prevention with a focus on vaccine evaluation.

## Ethics Statement

All the animal experiments were performed in accordance with the ethical guidelines of Huazhong Agricultural University (HZAU), and the protocols were approved by the Ethical Committee of HZAU (HZAUFI-2025-0061).

## Acknowledgements

This work was supported by the National Key Research and Development Program of China (2023YFD2402400, 2023YFD2400701), National Natural Science Foundation of China (32373162), Major Science and Technology Project of Hubei Province (2023BBA001), and HZAU-AGIS Cooperation Fund (SZYJY2023006). The authors sincerely thank Prof. Yu-Ding Fan, Yangtze River Fisheries Research Institute, Chinese Academy of Fishery Sciences, for generously providing the GCRV-Ⅱ (GCRV106), which served as a critical resource for this study and significantly facilitated our experimental progress.

## Notes

### Competing Interest Statement

The authors have declared no competing interest.

